# Comparative evaluation of straight and curved extension dialysis catheters for continuous renal replacement therapy in dogs with acute kidney injury

**DOI:** 10.1101/2023.06.11.544513

**Authors:** Abid Ali Bhat, M. Chandrasekar, A. P. Nambi, Sandhya Bhavani, S. Kavitha, Firdous A. Khan

## Abstract

A patent dual lumen dialysis catheter is one of the basic requirements for efficient extracorporeal (EC) therapy. The objective of this study was to measure resistance to blood flow offered by straight and curved extension dual lumen dialysis catheters used for continuous renal replacement therapy (CRRT). Twenty dogs suffering from acute kidney injury (AKI) were subjected to CRRT. The dogs were allocated randomly to Group-I (Curved extension catheter, n=12) or Group-II (Straight extension catheter, n=8), based on the type of dual-lumen catheter used in CRRT. The catheter outflow and inflow pressures were recorded at blood pump speeds of 50 mL/min and 99-100 mL/min. Data were tested for normality and differences in mean inflow and outflow catheter resistances were evaluated for statistical significance using independent samples *t* tests. Straight extension catheters offered lower inflow resistance than curved extension catheters at both 50 mL/min (41.50 ± 5.84 mm Hg and 63.75 ± 6.88 mm Hg, respectively; *P*=0.03) and 99-100 mL/min (63.00 ± 8.11 mm Hg and 86.92 ± 7.02 mm Hg, respectively; *P*=0.04) blood flow rates. Straight extension catheters also offered lower outflow resistance than curved catheters at 99-100 mL/min (−94.12 ± 7.91 mm Hg and -128.25 ± 7.56 mm Hg, respectively; *P*=0.01; the negative signs only indicate the direction of blood flow). These findings suggest that straight extension dual lumen dialysis catheters would likely perform better than the curved model in extracorporeal renal replacement therapy.

## Introduction

Acute kidney injury (AKI) is generally characterized by an abrupt reduction in kidney function and/ or reduced urine output. It is an ongoing process resulting in reduced functional mass^1^ and may or may not lead to renal failure.^2^ Due to the dearth of renal specific therapeutic agents, advancement in the management of AKI has been in the area of renal replacement therapy (RRT). Renal replacement therapies can effectively restore electrolyte, acid-base, and fluid balance, and eliminate retained uremic toxins, thereby prolonging survival and improving the potential for recovery.^3^ Vascular access is one of the primary requirements of successful RRT^4^ and its adequate patency allows smooth management of extracorporeal (EC) blood circuit.^5^ Appropriate functioning of the catheter during CRRT allows effective and adequate urea clearance. In animal practice, central venous catheters are the traditional form of vascular access^4^ and the jugular vein is the established site preferable in small animals. Understanding the facts about resistance to the blood flow shown by catheters with different designs may help clinicians in choosing an appropriate catheter and avert vascular access dysfunction. To the authors’ knowledge, there are no previous reports on the evaluation of resistance to blood flow by different types of catheters during CRRT in dogs. Therefore, this study aimed to determine resistance offered by straight and curved extension dual lumen dialysis catheters during continuous renal replacement therapy (CRRT) in dogs.

## Materials and Methods

### Animal Selection

This study was approved by the institutional animal care and use committee. Written informed consent was obtained from each owner before conducting the procedures.

Clinical cases confirmed as suffering from AKI were classified based on International Renal Interest Society classification (IRIS), 2013. Twenty dogs (Grade IV, n=5; Grade V, n=15) were subjected to CRRT and allocated randomly to Group-I (Curved extension catheter, n=12) and Group-II (Straight extension catheter, n=8) based on the type of Quinton-Mohurkar dual-lumen catheter used in CRRT.

### CRRT Procedure

An 11.5 Fr × 13.5 centimeter Quinton-Mohurkar dual-lumen dialysis straight catheter (with straight and curved extension) was placed into the left jugular vein (Fig.1) of dogs by Seldinger technique^6^ using a J-guide wire and venodilator under local anesthesia (Lidocaine 2 percent). No sedatives were given during the study period. The catheter was advanced down the vein until the tip was located close to the level of the right atrium. Each vascath lumen of the catheter was heparin locked (ampoule 5000 units in 5 ml) to prevent clotting. Heparin locking solution was aspirated before use. The catheter placement was confirmed by digital radiography in each patient. The catheter was secured by stay sutures and the cutaneous site was treated with povidone-iodine followed by protective bandage during the CRRT and adhesive elastic bandage on termination of the treatment.

**Fig.1.**
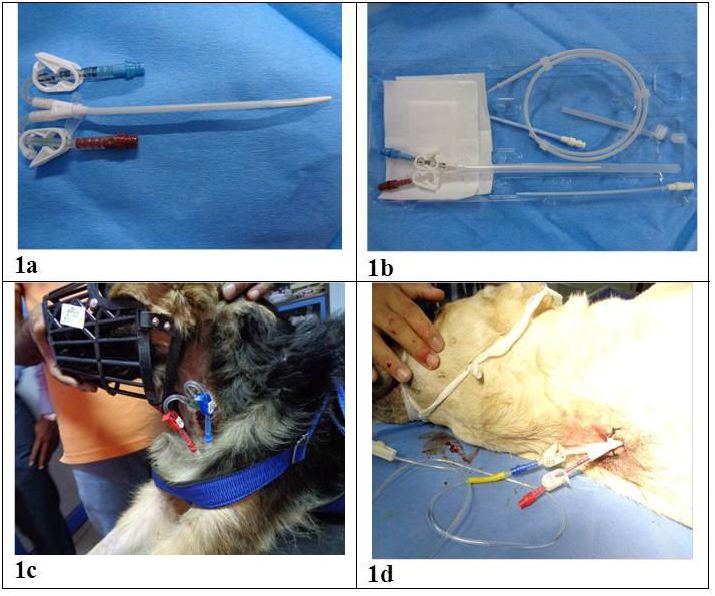
Quinton-Mohurkar Catheter Placement by Seldinger Technique 1a. Curved Catheter 1b. Straight Catheter 1c. Curved Catheter Placement by Seldinger Technique 1d. Straight Catheter Placement by Seldinger Technique

The catheter outflow (Ro) and inflow (Ri) pressures (resistance) were recorded a blood flow speed (QB) of 50 mL/min following 1 minute of blood flow stabilization after the patient was connected to the circuit. After the pressure readings were recorded at 50 mL/min, the blood flow speed was maintained for 5 minutes before switching over to 99-100 mL/min. The outflow and infow pressures were recorded from the in-blood-line pressure transducers built into the Prisma machine (Gambro Prismaflex, Gambro Lundia AB, SE-22010, Lund Sweden). At least five readings were recorded at each blood flow during the treatment and an average of the readings was used for analysis. The catheters were evaluated on the first treatment for all patients. The proximal lumen of the catheter was used as an inlet for all the patients. The treatment time ranged between 8 hours and 22 hours.

### Selection of Treatment Mode

Continuous venovenous hemodiafiltration (CVVHDF) treatment mode with postdialyzer configuration for replacement fluid was used in this study. Anticoagulation: The PrismaFlex CRRT circuit was primed with 500 mL of 0.9 % normal saline containing 5000 units of heparin. This facilitates heparin bonding to the filter membrane that is necessary to prevent filter membrane clotting. During CRRT, patients were managed as per the Louisiana State University heparin work sheet^7^.

### Statistical Analysis

Data were analyzed using SPSS statistics 17.0 software (SPSS Inc., Chicago, IL). Shapiro-Wilk and Kolmogorov-Smirnov tests were used to evaluate normality. As the data were normally distributed (*P*>0.05), differences in inflow and outflow resistance between straight and curved extension catheters at the two specified blood flow rates were evaluated for statistical significance by ‘independent samples *t* test’. Differences with *P*-values less than 0.05 were considered to be statistically significant. Results are presented as mean ± standard error (mean±SE).

## Results

The mean inflow resistance offered by straight extension catheters was lower than the resistance offered by curved extension catheters at 50 mL/min (41.50 ± 5.84 mm Hg and 63.75 ± 6.88 mm Hg, respectively; *P*=0.03) and 99-100 mL/min (63.00 ± 8.11 mm Hg and 86.92 ± 7.02 mm Hg, respectively; *P*=0.04) blood flow rates (Fig. 2a). Straight extension catheters also offered lower outflow resistance than curved extension catheters at 99-100 mL/min (−94.12 ± 7.91 mm Hg and -128.25 ± 7.56 mm Hg, respectively; *P*=0.01; the negative signs only indicate the direction of blood flow). A similar numerical trend was observed at 50 mL/min (−69.00 ± 6.68 mm Hg and -82.08 ± 6.78 mm Hg, respectively; *P*=0.21), however, the difference was not statistically significant (Fig. 2b). The inflow and outflow resistances for both catheters showed a consistent rise as the blood flow rate was increased from 50 mL/min to 99-100 mL/min.

**Fig. 2.**
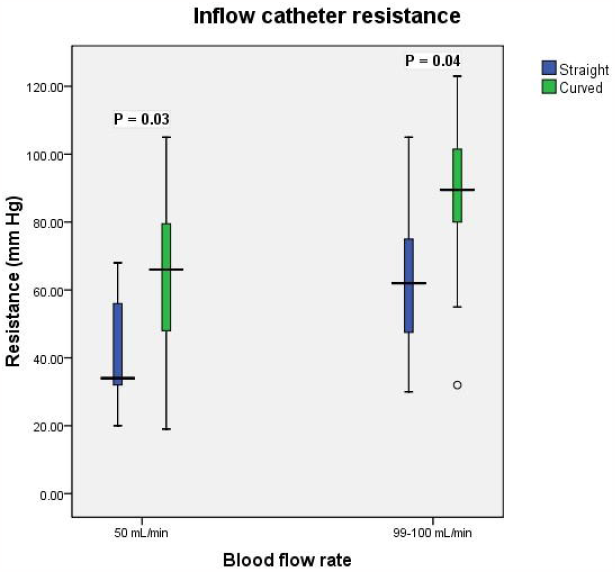
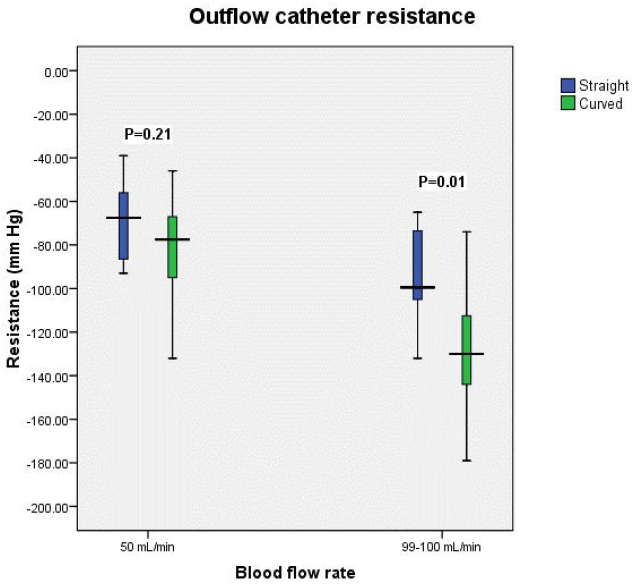
Box and whisker plots showing inflow catheter resistance (2a) and outflow catheter resistance (2b) offered by straight and curved catheters. Within each blood flow rate, *P*-values less than 0.05 indicate a significant difference in mean resistance between the two groups. By convention, inflow catheter resistance is expressed as positive and outflow catheter resistance is expressed as negative. The positive and negative signs only indicate the direction of blood flow and not the actual measurements.

## Discussion

Continuous renal replacement therapy is commonly used in critically ill human patients with acute kidney injury.^8,9^ However, due to lack of facility and expertise, CRRT is not widely used in small animal practice. The efficacy of CRRT depends on the continuous patency of the extracorporeal circuit (EC). Patent vascular access is a major prerequisite for successful treatment delivery and adequacy. Clot formation in the catheter, however, remains a major concern that reduces CRRT adequacy, access lifespan, and results in frequent circuit breakdown. Moreover, continuously patent catheter allows for smooth and efficient patient management. Prolongation of a continuously patent EC is of central importance and is usually accomplished by anticoagulation. An increase in systemic/circuit anticoagulation dose is a common response to frequent EC clotting. However, this methodology leads to increased risk of bleeding and ignoring the factors related to inadequate vascular access that may be of similar importance in the pathogenesis of circuit clotting.^5^

In our study, both the catheters were associated with different blood flow resistances over the specified blood flow rate which is in congruence with Tan et al.^5^ Straight extension catheters offered lower resistance as compared to curved extension catheters and proved to be an ideal choice for CRRT. Higher blood flow resistance may result in access pressure alarms indicating a failure to maintain EC blood flow. In a human study, catheter dysfunction either due to infectious or noninfectious complications leads to increased mortality in the affected population.^10^ However, such studies are lacking in veterinary medicine. Moreover, catheter function can decrease with time if thrombosis or stenosis develops gradually.^4^

In small animal practice, acute kidney injury cases are usually hemodynamically unstable due to dysregulation of fluid, electrolytes, and acid-base imbalance^11^. Catheter malfunction is a major concern in such animals undergoing renal replacement therapy. There are several hemodynamic factors that promote thrombosis and stasis of blood flow. These factors include restenosis due to surgical manipulation, vessel shear strain, pressure, and vascular remodeling in response to injury.^12^ Vascular endothelial cells can sense and respond to changes in the forces acting on them especially oscillatory pressure changes by modulating expression at the cellular level.^12^ Moreover, this response may be aggravated in the presence of high resistance offered by the dual-lumen catheters causing oscillatory pressure changes leading to perpetuating endothelial pathological response. Therefore, a vicious cycle sets in, which corrodes the adequacy of CRRT^5^. These factors are usually unavoidable; however, the type and design of catheters chosen can be manipulated and might help in reducing several stressors impairing blood flow. This notion appears to be supported by the present study demonstrating that catheter performance measured as inflow and outflow resistance to blood during CRRT varies according to the design of the catheter. Further studies are required to investigate the endothelial changes associated with use of different catheters during CRRT in dogs.

The present study has some limitations that should be considered while interpreting or applying its findings. Since a completely randomized design was followed for allocation of the patients to the two study groups (curved and straight extension catheters), there was an unequal sample size in the two groups. A strength of this approach though is that it minimizes selection bias. Another limitation of the study was the use of a single length catheter in all dogs due to non-availability of different length catheters. However, the study involved small and medium sized dogs and catheters were inserted in the middle to lower third of the jugular vein to ensure that the catheter tip was close to the level of right atrium. Nevertheless, the findings should be interpreted with caution, considering a potential confounding effect of catheter length in dogs of different sizes. The impact on comparison between the straight and curved extension catheter groups is likely minimal though because the dogs were allocated randomly to the two groups. The small sample size of this study is also a limitation and, therefore, the findings need to be confirmed in further studies involving larger populations of AKI-affected dogs undergoing CRRT. Finally, catheters were placed in the left jugular vein in all of the dogs used in this study due to the attending clinicians’ preference for the left side. Although the catheter placement on the same side in all the dogs minimizes confounding, selection of the left side limits the potential extrapolation of results to a general canine population in which the right jugular vein is catheterized more often than the left jugular vein. Further studies involving more optimal placement of the catheters are required to confirm the preliminary findings of this study.

## Conclusion

In conclusion, results of the current study indicate that straight extension Quinton-Mohurkar dialysis catheters offer lower outflow and inflow resistance than the curved extension model in dogs undergoing CRRT. Further studies are required to confirm these findings in a broader population..

## Acknowledgments

The authors are thankful to the dog owners for their cooperation and the supporting staff members for their assistance.

## Authors’ Contributions

AAB and APN conceived the study; AAB and MC performed the animal procedures; FAK analyzed the data; AAB and FAK wrote the manuscript; SB and SK critically reviewed the manuscript. All authors read and approved the final manuscript.

## References

1. Mehta RL, Sudhir JA, Shah V, et al. Acute Kidney Injury Network: report of an initiative to improve outcomes in acute kidney injury. Crit Care; 11: R31, 2007

2. Thoen ME, Kerl ME. Characterization of acute kidney injury in hospitalized dogs and evaluation of a veterinary acute kidney injury staging system. J Vet Emerg Crit Care 21: 648–657, 2011

3. Sagev G, Palm C, LeRoy B, et al. Evaluation of neutrophil gelatinase-associated lipocalin as a marker of kidney injury in dogs. J Vet Intern Med; 27: 1362–136, 2013

4. Chalhoub S, Langston CA, Poeppel K. Vascular access for extracorporeal replacement therapy in veterinary patients. Vet Clin North Am Small Anim Pract 41: 147–161, 2011

5. Tan HK, Bridge NI Baldwin, et al. Ex-vivo evaluation of vascular catheters for continuous hemofiltration. Ren Fail 24: 755–762, 2002

6. Davenport A. Anticoagulation for continuous renal replacement therapy. Contrib Nephrol 144: 228–38, 2004

7. Acierno MJ. Continuous Renal Replacement therapy in dogs and cats. Vet Clin North Am Small Anim Pract 41: 135–146, 2011

8. Ronco C, Cruz D, Bellomo R. Continuous renal replacement therapy in critical illness. Contrib Nephrol 156: 309–319, 2007

9. Kim I, Fealy N, Baldwin I, et al. A comparison of the Niagara™ and Dolphin^®^ catheters for continuous renal replacement therapy. Int J Artif Organs 34: 1061–1066, 2011

10. Miller LM, Clark E, Dipchand C, et al. Hemodialysis Tunneled Catheter-Related Infections. Can J Kidney Health Dis 3: 1–11, 2016

11. Bloom CA, Labato MA. Intermittent hemodialysis for small animals. Vet Clin North Am Small Anim Pract 41: 115–133, 2011

12. Walsh P, Mclachlan C. Stenosis and thrombosis. In: Metin Akay, ed. Wiley Encyclopedia of Biomedical Engineering. New Jersey, USA: John Wiley & Sons, 2006:1–12.

